# Profiling of pluripotency factors in individual stem cells and early embryos

**DOI:** 10.1101/286351

**Authors:** Sarah J. Hainer, Ana Bošković, Oliver J. Rando, Thomas G. Fazzio

## Abstract

**Major cell fate decisions are governed by sequence-specific transcription factors (TFs) that act in small cell populations within developing embryos. To understand how TFs regulate cell fate it is important to identify their genomic binding sites in these populations. However, current methods cannot profile TFs genome-wide at or near the single cell level. Here we adapt the CUT&RUN method to profile chromatin proteins in low cell numbers, mapping TF-DNA interactions in single cells and individual pre-implantation embryos for the first time. Using this method, we demonstrate that the pluripotency TF NANOG is significantly more dependent on the SWI/SNF family ATPase BRG1 for association with its genomic targets *in vivo* than in cultured cells—a finding that could not have been made using traditional approaches. Ultra-low input CUT&RUN (uliCUT&RUN) enables interrogation of TF binding from low cell numbers, with broad applicability to rare cell populations of importance in development or disease.**

## INTRODUCTION

Cellular heterogeneity presents a significant obstacle to the study of complex systems in metazoans (Yuan et al., 2017). Key developmental processes are often initiated in small populations of cells that expand and differentiate to generate complex tissues within the embryo. In adults, rare tissue-specific stem cells act in response to stimuli or damage to maintain tissue homeostasis. In addition, cells in some cancer types with properties of stem cells facilitate regeneration of the tumor mass after therapy. Because of the important roles of rare stem and progenitor cell populations in each of these settings, sensitive methods for characterizing their regulation and functions are necessary to better understand development and disease.

Cell fate decisions are orchestrated in large part by the concerted actions of TFs and chromatin remodeling proteins. Expression of lineage-specific TFs leads to the activation and repression of specific sets of genes that dictate cell identity, while chromatin remodeling proteins facilitate and help enforce changes in gene expression (Young, 2011). The functions of developmental TFs and chromatin remodeling enzymes are interdependent—while some TFs direct chromatin remodeling proteins to specific regulatory regions, chromatin remodeling at enhancers is necessary for binding of other TFs with roles in directing cell fate (Zaret and Mango, 2016). Accordingly, comprehensive maps of the binding sites of TFs and chromatin regulators are necessary to understand how gene expression patterns are rewired during cell fate changes.

Unfortunately, current methods for mapping the genomic locations of TFs and chromatin remodeling enzymes are insufficiently sensitive to allow mapping in small populations of cells. As an alternative strategy, methods for identification of “open” chromatin regions have enabled inference of regulatory elements such as enhancer and silencer elements, which are generally accessible to non-sequence-specific enzymes that generate DNA breaks (Buenrostro et al., 2013; Crawford et al., 2004; Sabo et al., 2004). Highly sensitive modifications of two such methods, ATAC-seq and DNase-seq, have recently enabled analysis of chromatin accessibility in single cells, allowing examination of epigenomic variability and identification of cell type-specific chromatin features (Buenrostro et al., 2015b; Cusanovich et al., 2015; Jin et al., 2015). While these techniques are extremely powerful for discovery of regulatory features, in many cases it is not possible to identify the regulatory proteins that render chromatin structure accessible at each site. Chromatin remodeling enzymes generally bind without preference for DNA sequence, preventing identification of DNA sequence motifs specific for these factors within open chromatin regions. Even for sequence-specific TFs, many enhancers include binding sites for multiple TFs. Furthermore, groups of related TFs often bind similar motifs. Therefore, high confidence assignment of accessible chromatin peaks to any one TF is not possible in most cases.

Chromatin immunoprecipitation (ChIP) is a widely used technique for exploring protein-DNA interactions on chromatin. Modifications to traditional ChIP protocols have been developed in recent years to profile TF binding in small populations of cells, including ChIPmentation, carrier-assisted ChIP-seq, ULI-NChIP, μChIP, and DROP-ChIP (Brind’Amour et al., 2015; Dahl et al., 2016; X. Liu et al., 2016; Rotem et al., 2015; Schmidl et al., 2015). Several of these techniques enable mapping of abundant histone modifications such as H3K4me3 and H3K27me3 in fewer than 1,000 cells. However, ChIP-based methods for mapping chromatin occupancy of TFs currently require 10,000 cells or more to observe reproducible peaks of enrichment (Schmidl et al., 2015; Zwart et al., 2013). An alternative approach utilizing overexpression of bacterial Dam methylase fused to TFs allows profiling from as few as 1,000 cells (Tosti et al., 2017). However, overexpression of fusion proteins may not be feasible in some settings and could potentially lead to occupancy at non-physiological locations. Because of these difficulties, the genomic landscape of TF binding in single cells, individual preimplantation embryos, or other rare populations has been thus far inaccessible.

CUT&RUN is a recently described method for genome-scale profiling derived from ChIC, in which a recombinant protein A-micrococcal nuclease (MNase) fusion protein is recruited via antibodies to the genomic locations of chromatin proteins, and underlying DNA fragments are liberated from bulk chromatin by endonucleolytic cleavage (Schmid et al., 2004; Skene and Henikoff, 2017a). CUT&RUN has a number of advantages over traditional ChIP-based techniques, most importantly specific DNA digestion by targeted MNase results in low background leading to increased enrichment and a decreased requirement for high read coverage. CUT&RUN has been successfully used to map H3K27me3 genome-wide using as few as 100 cells and the insulator protein CTCF from as few as 1,000 (Skene and Henikoff, 2017b).

Despite these advances, there are still no methods capable of profiling TF occupancy genome-wide in single cells or individual embryos. In this study, we adapt the CUT&RUN method for ultra-low input by modifying several steps in the original protocol. These modifications enable us to profile the genomic occupancies of several chromatin proteins in small populations of cells, individual pre-implantation mouse embryos, and even in single cells. Using this technique, we then test the extent to which properties of TF binding previously measured in cell culture are shared *in vivo*—a question that could not be addressed using traditional mapping approaches. We focused on the pluripotency TF NANOG, which has previously been shown to display minimal requirement for the SWI/SNF family ATPase BRG1 for binding to its genomic targets in cultured mouse embryonic stem cells (mESCs) (King and Klose, 2017). Interestingly, we find that NANOG is highly dependent on BRG1 for association with its genomic targets in mouse blastocysts, suggesting NANOG is significantly more sensitive to the underlying chromatin environment for association with its targets *in vivo*. Together, we show that uliCUT&RUN is able to profile the occupancy of DNA binding proteins from extremely small populations, permitting the study of TF binding in biologically relevant populations *in vivo* that are difficult to obtain in large numbers.

## RESULTS AND DISCUSSION

### Adaptation of CUT&RUN for very low cell numbers

In order to profile chromatin proteins from fewer than 1,000 cells, we altered the original CUT&RUN protocol to optimize for ultra-low input (see methods); we denote the modified protocol as “ultra-low input CUT&RUN” (uliCUT&RUN). We used uliCUT&RUN to profile occupancy of the insulator protein CTCF and of histone H3 lysine 4 tri-methylation (H3K4me3), a mark of active promoter regions, from populations of mESCs ranging in number from 500,000 to 10. Based on the different size distributions of MNase footprints for sequence-specific binding factors and nucleosomes (Skene and Henikoff, 2017a), we selected reads of less than 120 bp in length after paired-end sequencing of CTCF libraries and 150-500 bp reads for H3K4me3 (Figure S1A-B). Focusing on the genomic locations of CTCF and H3K4me3 previously identified by ChIP-seq from millions of mESCs—taken as the “gold standard” here—we observed significant enrichment of CTCF and H3K4me3 relative to surrounding regions at each cell number tested (Figure 1A-B and S1C-D). In contrast, control samples lacking a primary antibody (referred to as “no antibody”) exhibited minimal enrichment over the same regions. Nucleosome-sized reads from CTCF uliCUT&RUN maps revealed the expected pattern of well-positioned nucleosomes immediately flanking CTCF binding sites (Figure S1E), further validating these data. Over a broad range of cell numbers, we found that peaks of uliCUT&RUN enrichment strongly overlapped with peaks from high input ChIP-seq data (Figure 1C-D and S1F-G). Consistent with these findings, 10, 50, and 500 cell maps of both CTCF and H3K4me3 were very similar to high cell number maps at numerous genomic locations (examples shown in Figure 1E-F). Collectively, these findings demonstrate consistent and sensitive mapping of two distinct chromatin proteins from low numbers of cells using uliCUT&RUN.

**Figure 1.**
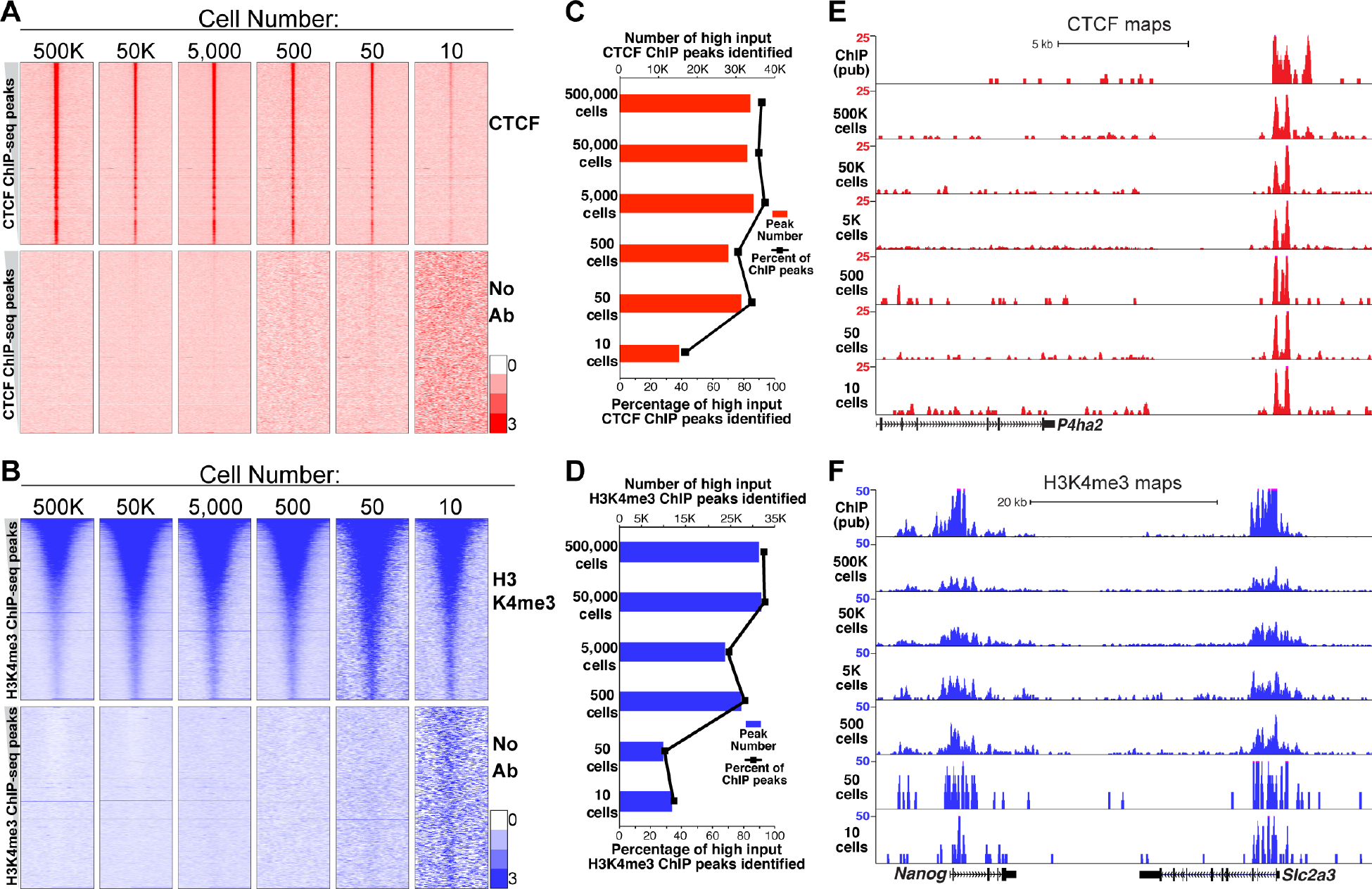
Localization of chromatin proteins from low cell numbers using uliCUT&RUN. **(A)** uliCUT&RUN data for CTCF and no antibody (No Ab) for indicated cell numbers. Heatmap rows correspond to normalized read density surrounding CTCF binding sites (center) called from GSE11431 with 2 kb of adjacent sequence on each side. Rows are ranked from highest ChIP-seq enrichment (top) to lowest (bottom). **(B)** uliCUT&RUN data for H3K4me3. Data are centered on peaks called from GSE31039 and organized as in (A). **(C-D)** Number and percentage of high input ChIP-seq peaks identified by uliCUT&RUN for CTCF (C) or H3K4me3 (D) from indicated cell numbers. **(E-F)** Browser tracks comparing normalized read enrichment at sites of CTCF (E) or H3K4me3 (F) enrichment.

### Robust profiling of pluripotency TFs from 50 cells by uliCUT&RUN

CTCF and H3K4me3 have proven to be among the most robust epitopes for ChIP-based studies, raising the question of whether uliCUT&RUN can effectively map a broader array of DNA-binding factors. To explore the general utility of uliCUT&RUN for mapping diverse DNA-binding proteins, we generated 50,000 and 50 cell profiles for several TFs, histone modifications, and a nucleosome remodeling enzyme in mESCs. In each case, uliCUT&RUN profiles from both 50 and 50,000 cells showed enrichment at genomic locations previously determined by high input ChIP-seq (Figure 2A-I and S2A-C). We note that even in control experiments lacking a primary antibody, TF binding sites exhibited a subtle aggregate enrichment relative to surrounding regions, consistent with the fact TF-bound regions are hypersensitive to nucleases, including the untargeted protein A-MNase used in these controls (Figure 2G-I). However, the no antibody controls exhibited minimal correlation with ChIP-seq enrichment at the level of individual genes (Figure 2A-F). These findings confirm the specificity of uliCUT&RUN maps for each factor tested, in both 50,000 and 50 cell samples.

**Figure 2.**
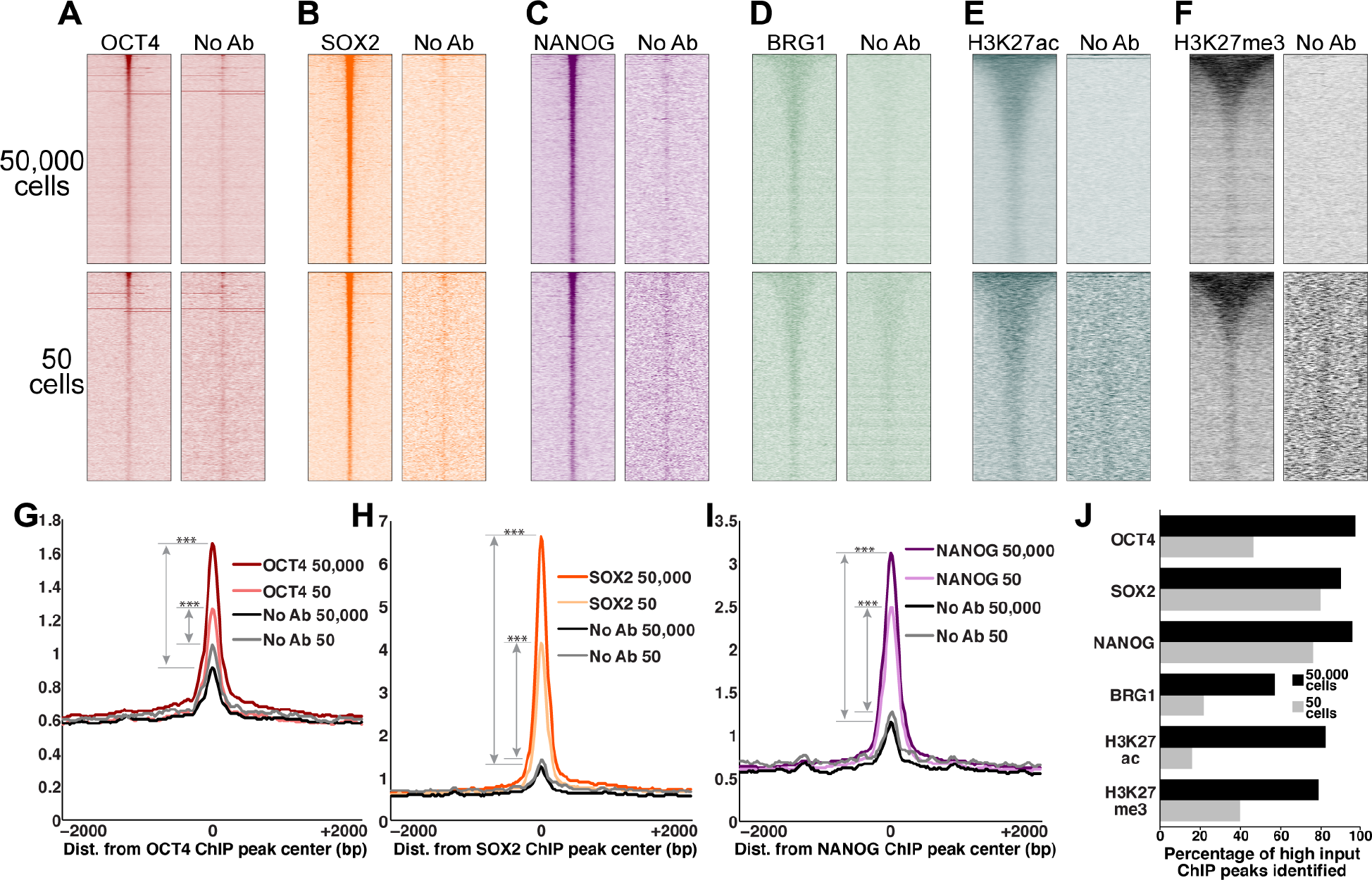
Sensitive mapping of TFs and chromatin proteins using uliCUT&RUN. **(A-F)** uliCUT&RUN enrichment of indicated chromatin proteins from 50 and 50,000 mESCs. Heatmaps are organized as in Figure 1, depicting uliCUT&RUN enrichment at ChIP-seq binding sites for OCT4 (GSE11724, A); SOX2 (GSE11724, B); NANOG (GSE11724, C); BRG1 (GSE14344, D); H3K27ac (GSE31039, E); and EZH2 (GSE49435, F). **(G-I)** Average enrichment over all binding sites of OCT4 (G), SOX2 (H), or NANOG (I). ***p<2.2×10^−16^ (K-S test; see methods for details). **(J)** Percentage of high input ChIP-seq peaks identified by uliCUT&RUN using 50 or 50,000 cells.

Next, we compared peaks of uliCUT&RUN enrichment for each factor obtained from low and high cell numbers. For TFs SOX2 and NANOG, the majority (76-96%) of “gold standard” binding sites were identified in both the 50 and 50,000 cell samples, and approximately half (47%) of OCT4 binding sites were identified from 50 cell uliCUT&RUN profiling (Figure 2J). Maps of the SWI/SNF ATPase Brg1, as well as two well-studied histone modifications, H3K27me3 and H3K27ac, revealed moderate overlap with established binding sites in 50 cell samples (Figure 2J). Furthermore, for all factors examined, 50 and 50,000 cell peaks were highly overlapping with each other (Figure S2D). Finally, the established DNA sequence motifs corresponding to OCT4, SOX2, and NANOG were significantly enriched within the 50 and 50,000 cell uliCUT&RUN peaks corresponding to each factor (Figure S2E), further demonstrating the specificity of uliCUT&RUN TF maps. Collectively, these data demonstrate an increase in sensitivity of at least three orders of magnitude for TF mapping relative to ChIP-based approaches.

### Single cell TF profiling using uliCUT&RUN

In principle, the capacity of uliCUT&RUN to profile TF occupancy from 10-50 cells enables interrogation of even the most limited biological samples, such as preimplantation embryos or small tissue specimens. However, single cell experiments can capture features of gene regulation missed in studies of cell populations (Z. Liu and Tjian, 2018; Stubbington et al., 2017). We therefore examined the feasibility of single cell TF mapping experiments using uliCUT&RUN. To this end, we sorted single mESCs into individual wells of a 96-well plate and performed uliCUT&RUN with antibodies specific for CTCF, SOX2, or NANOG (see methods). Initial studies at low sequencing depth showed variable levels of enrichment across individual cells (Supplementary Table 1 and Figure 3A). Nonetheless, deeper sequencing revealed that single-cell libraries captured overall enrichment of all three factors at their established binding sites (Figure 3B-D).

**Figure 3.**
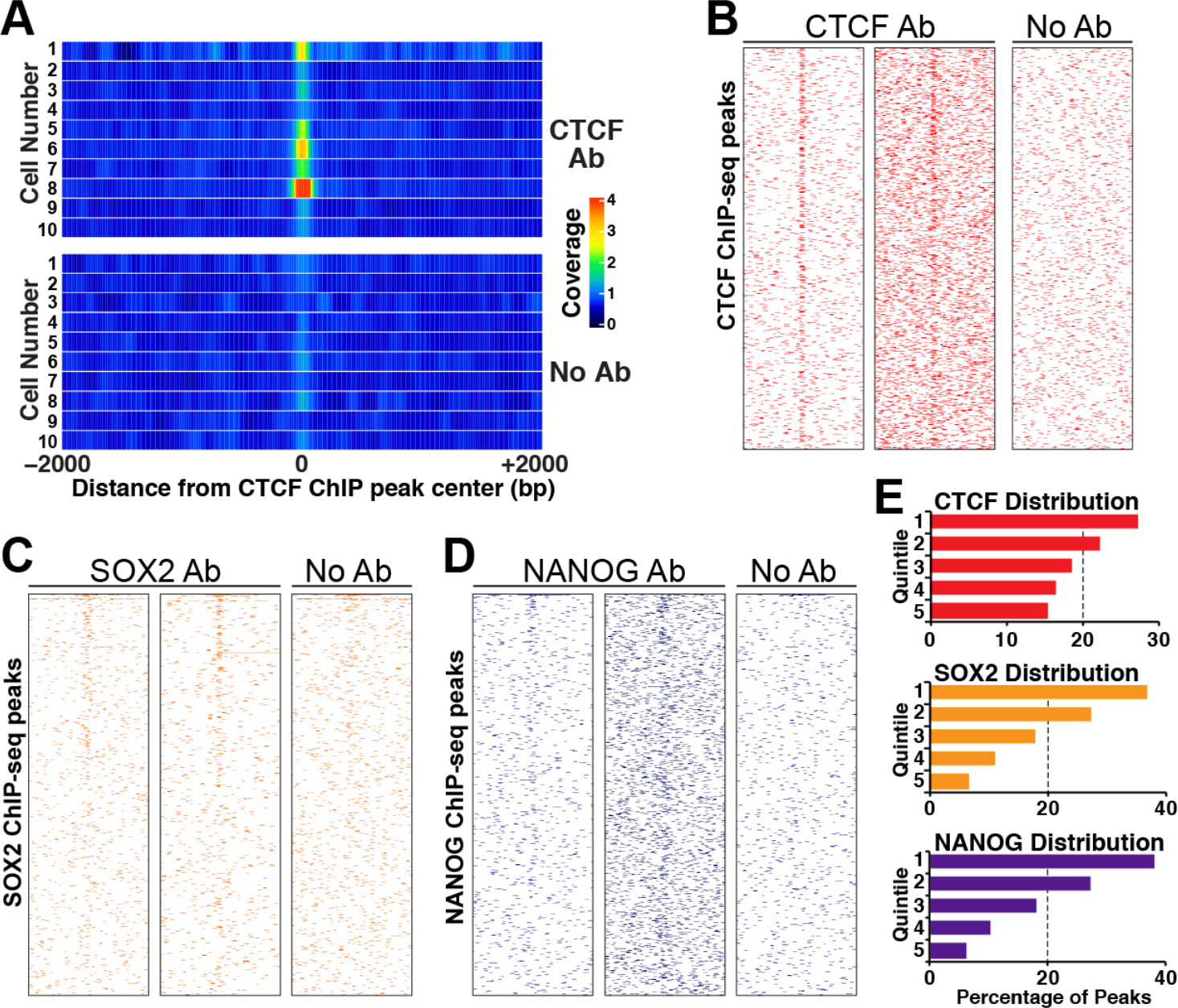
Single cell TF mapping by uliCUT&RUN. **(A)** Enrichment at known CTCF binding sites for single cells subjected to uliCUT&RUN. For each cell, average enrichment over CTCF binding sites is shown as a one-dimensional heatmap. **(B-D)** Heatmaps depicting single cell uliCUT&RUN data for CTCF (B), SOX2 (C), or NANOG (D). ChIP binding sites with no uliCUT&RUN read coverage within the 4 kb window are not shown. **(E)** The distribution of single cell uliCUT&RUN peaks within five high input ChIP-seq quintiles from most enriched (1) to least (5) are shown.

As with single cell ATAC-seq and DNase-seq, which capture only a small fraction of the accessible sites identified from large populations of cells (Buenrostro et al., 2015b; Cusanovich et al., 2015; Jin et al., 2015), single cell uliCUT&RUN captured a portion of TF binding sites from high input ChIP-seq maps (Supplementary Table 1). However, compared to the fraction of high input ATAC-seq peaks (Q. Liu et al., 2017) identified by single cell ATAC-seq (Buenrostro et al., 2015b), single cell uliCUT&RUN libraries identified “gold standard” TF binding sites at a higher rate on average (Figure S3), demonstrating uncommonly high sensitivity of single cell profiling using the uliCUT&RUN method. For both methods, the incomplete identification of high cell number peaks likely reflects both the technical challenge of recovering every fragment of DNA released from a single nucleus and meaningful biological variation. In particular, TF binding sites within peaks from high cell number maps are likely occupied in only a fraction of all cells in the population. Accordingly, weak to moderate peaks may represent TF binding in a small minority of cells.

Finally, to test whether single cell uliCUT&RUN libraries were more likely to identify the most highly occupied TF binding sites throughout the genome, we divided high input ChIP-seq peaks into quintiles based on their level of enrichment from most enriched (1^st^ quintile) to least (5^th^ quintile). We sorted each overlapping single cell uliCUT&RUN peak into the quintile containing its matching ChIP peak and found that single cell peaks were overrepresented among the top quintiles (Figure 3E), consistent with our expectations. Collectively, these findings reveal that uliCUT&RUN maps of single cells are accurate but necessarily incomplete representations of CTCF, SOX2, and NANOG binding in mESCs. Future applications of single cell uliCUT&RUN should uncover differences in TF occupancy between distinct cell types or different locations of cells *in vivo*, among other uses.

### Association of NANOG with its genomic targets is dependent on BRG1 *in vivo*

OCT4, SOX2, and KLF4, and NANOG are critical for pluripotency in inner cell mass cells of blastocyst stage embryos, as well as their cultured counterparts, ESCs (Young, 2011). In mESCs, BRG1—the catalytic component of the esBAF nucleosome remodeling complex that activates enhancers by creating open chromatin structure—is required for association of OCT4 with approximately 60% of its normal genomic binding sites (Hainer and Fazzio, 2015; Hodges et al., 2018; Hu et al., 2011; King and Klose, 2017). In contrast, chromatin association of SOX2 and NANOG is only modestly dependent on BRG1 function (Hainer and Fazzio, 2015; King and Klose, 2017), suggesting that continuous chromatin remodeling is dispensable for sustained binding of these factors at most loci. However, the *establishment* of pluripotency factor binding as the zygote develops into the blastocyst has not been investigated. BRG1 is maternally deposited in oocytes and functions starting at zygotic genome activation (Bultman, 2006), whereas NANOG is expressed only at late morula and blastocyst stages (Chambers et al., 2003; Mitsui et al., 2003). Therefore, although NANOG binding is largely unaffected in BRG1-depleted mESCs, BRG1 may be required to open chromatin structure prior to the blastocyst stage in order to allow initial NANOG binding. Such a possibility has not been addressed on a genome-wide level because blastocysts are composed of ~30-80 cells and are therefore poorly suited to ChIP-based approaches.

To test the possibility that BRG1 is critical for chromatin association of NANOG *in vivo*, we adapted uliCUT&RUN to map binding of factors in single blastocysts (see methods). Pilot experiments mapping localization of CTCF in individual blastocysts demonstrated reproducible enrichment at “gold-standard” CTCF binding sites identified in ESCs (Figure 4A-B). These initial experiments demonstrate that uliCUT&RUN can be adapted for use in single pre-implantation embryos. Next, we tested the effect of BRG1 (gene name: *Smarca4*) depletion on genome-wide association of NANOG. We injected one cell mouse embryos with previously validated endoribonuclease prepared siRNAs (esiRNAs) targeting *Smarca4, Nanog*, or *EGFP* (Fazzio et al., 2008; Hainer et al., 2015), cultured each embryo to the early blastocyst stage (~30-50 cells), and mapped NANOG enrichment using uliCUT&RUN. Knockdown (KD) of each factor was confirmed by RT-qPCR using the cytoplasmic fraction that is normally discarded during the CUT&RUN procedure, as well as immunostaining of a parallel set of embryos (Figure S4A-C). Consistent with previous studies (Carey et al., 2015), *Smarca4* KD had no observable effect on *Nanog* expression in the inner cell mass but caused a modest increase in *Nanog* expression in trophoblast cells (Figure 4C), resulting in moderately elevated *Nanog* levels overall (Figure S4A-B). Relative to *EGFP* KD, *Nanog* KD strongly reduced NANOG enrichment at its genomic binding sites, demonstrating the specificity of *in vivo* uliCUT&RUN (Figure 4D). Interestingly, *Smarca4* KD also caused a strong reduction of NANOG enrichment across the genome (Figure 4D-E). As a physiologically relevant example, we zoomed in on the distal enhancer of the *Nanog* gene, where NANOG has been shown to bind and regulate its own expression (Boyer et al., 2005; Levasseur et al., 2008; Loh et al., 2006). We observed NANOG enrichment at this site in all control blastocysts, whereas enrichment was low or undetectable in three of four *Smarca4* KD embryos (Figure 4F).

**Figure 4.**
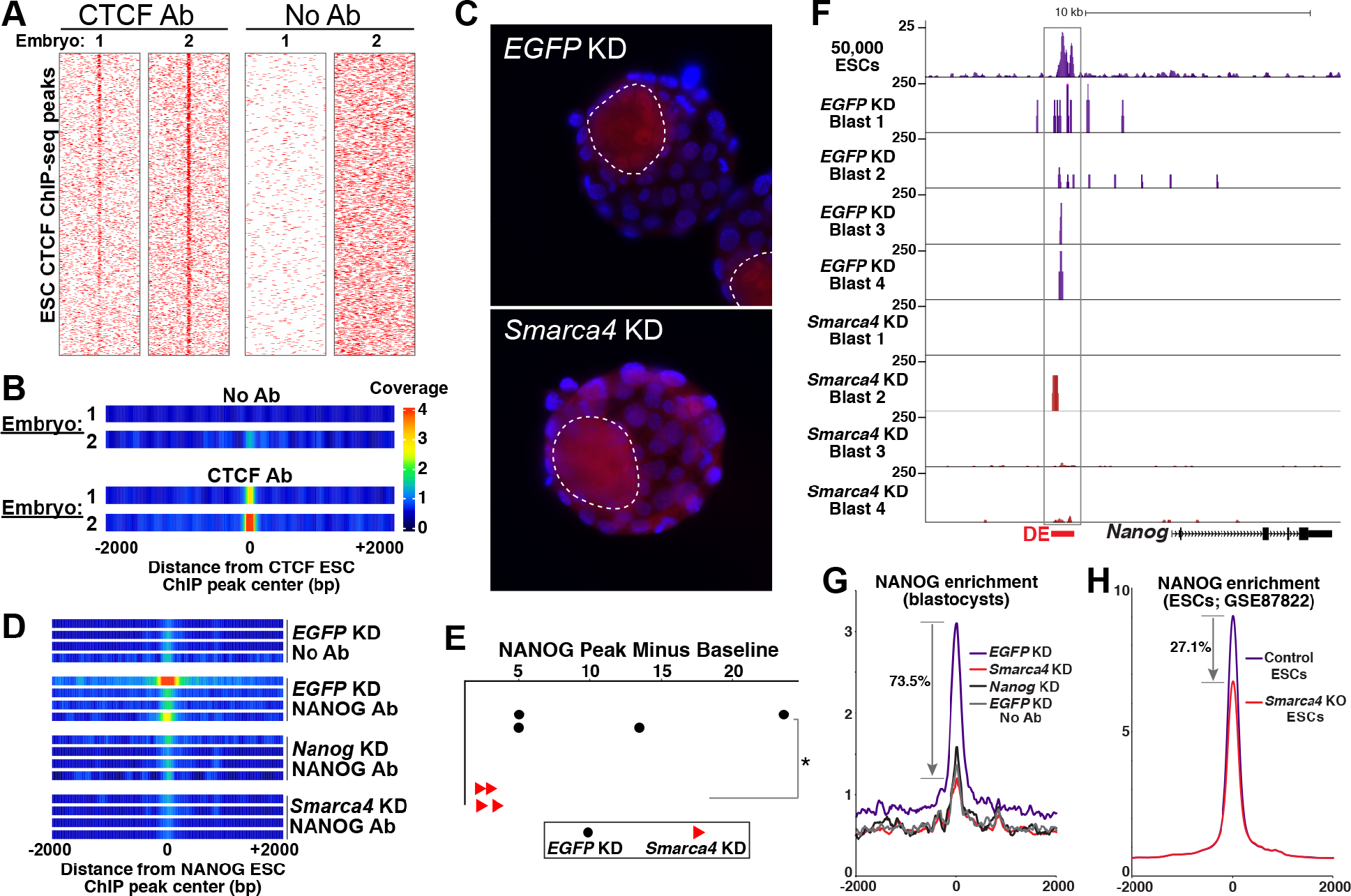
Chromatin association of NANOG is dependent on BRG1 *in vivo*. **(A)** CTCF or no antibody uliCUT&RUN maps of single blastocysts are shown as heatmaps sorted by high input CTCF ChlP-seq data from mESCs. **(B)** One dimensional heatmaps showing aggregate enrichment over CTCF binding sites. **(C)** NANOG immunofluorescence (red) of *EGFP* or *Smarca4* KD blastocysts. Boundaries of each inner cell mass are highlighted (dotted lines) and DAPI stained nuclei are shown in blue. **(D)** One dimensional heatmaps of *EGFP* or *Smarca4* KD embryos (four per group) subjected to uliCUT&RUN with NANOG antibody or no antibody. **(E)** Quantification of aggregate NANOG enrichment in embryos. Peak minus baseline is plotted for each embryo (see methods). *p<0.05; Mann-Whitney test. **(F)** Browser tracks showing NANOG enrichment in *EGFP* or *Smarca4* KD embryos (ESC data are shown for reference). The *Nanog* distal enhancer (DE) is highlighted. **(G-H)**, Changes in NANOG enrichment following *Smarca4* depletion in (G) blastocysts measured by uliCUT&RUN or (H) previously published ESC ChIP-seq data (King and Klose, 2017). Blastocyst replicates corresponding to each KD were averaged, as were ESC replicates corresponding to each KO.

We next quantified the extent to which NANOG binding depends on BRG1 in blastocysts and mESCs. In aggregate, we observed a 73.5% average reduction of NANOG enrichment—almost matching the enrichment observed in nonspecific control maps—in blastocysts following *Smarca4* KD (Figure 4G). In contrast, analysis of published ChIP-seq data from *Smarca4* knockout mESCs (King and Klose, 2017) revealed 27.1% aggregate reduction (Figure 4H), demonstrating that NANOG association with its target sites is much more dependent on BRG1 function *in vivo* than in mESCs. Critically, previous studies showed that depletion of *Smarca4* did not disrupt development to the blastocyst stage (Bultman et al., 2000; Kidder et al., 2009) and we observed no developmental delays or morphological changes upon *Smarca4* KD, ruling out this potential confounding factor. However, given the important roles for BRG1 in regulation of gene expression, potential indirect effects of *Smarca4* KD on NANOG localization cannot be ruled out.

In summary, we have shown that uliCUT&RUN is a powerful method for mapping the genomic locations of chromatin proteins, allowing for the first time the mapping of TFs from single cells and individual pre-implantation embryos. The increase in sensitivity for TF mapping is at least 1,000-fold relative to ChIP-based approaches. Using uliCUT&RUN, we demonstrate that NANOG is significantly more dependent on esBAF for chromatin association *in vivo* than in ESCs—a result that would not have been attainable using conventional mapping methods. This raises the possibility that other developmentally important TFs may have different requirements for chromatin association in embryos than has been observed in cell culture studies. We conclude that uliCUT&RUN is a powerful tool for interrogation of TFs in limited populations of cells, with high potential applicability in many additional *in vivo* contexts.

## ACKNOWLEDGEMENTS

We thank S. Henikoff for generously providing purified pA-MN protein and helpful advice on the CUT&RUN method, and the UMMS Flow Cytometry core for assistance. We thank J. Benanti, I. Bach, J. Dekker, and M. Walhout for critical reading of the manuscript. This work was supported by NIH grants R01HD072122 (to T.G.F.) and R01HD080224 (to O.J.R.). A.B. is supported by a fellowship from the Human Frontier Science Programme (LT000857/2015-L). S.J.H. is a Special Fellow of the Leukemia and Lymphoma Society. T.G.F. is a Leukemia and Lymphoma Society Scholar.

## AUTHOR CONTRIBUTIONS

S.J.H., A.B., O.J.R., and T.G.F. designed the study, wrote, and edited the manuscript. S.J.H. performed most experiments. A.B. performed embryo knockdown, culture, and immunofluorescence experiments. S.J.H. analyzed the data with assistance from T.G.F.

## METHODS

### Cell culture

E14 mouse ES cells (Hooper et al., 1987) were cultured as previously described (Chen et al., 2013). Cells have been verified that they are of male mouse origin through sequencing performed in this and previous studies and were previously tested to ensure they were free of mycoplasma.

### Antibodies

Antibodies used in this study were H3K4me3 (Millipore 05-745R), H3K27ac (Abcam ab4729), H3K27me3 (Millipore 07-449), CTCF (Millipore 07-729), OCT4 (Thermofisher 701756), SOX2 (Active Motif 39843), NANOG (Active Motif 61419), and BRG1 (Bethyl Labs A300-813).

### uliCUT&RUN procedure and library preparation

*Nuclei prep:* The CUT&RUN protocol was modified from Skene and Henikoff (Skene and Henikoff, 2017a); their detailed protocol is available online (http://blocks.fhcrc.org/steveh/papers/CUT&RUN_protocol.htm). Mouse ES cells were counted using a TC-10 cell counter (Biorad) and diluted to respective cell amounts. Cells were pelleted at 600g for 3 minutes at 4°C, the supernatant was discard and cells were washed with 1mL cold PBS. Cells were pelleted at 600g for 3 minutes at 4°C, the supernatant was discard and cells were resuspended in 1mL cold nuclear extraction (NE) buffer (20mM HEPES-KOH, pH 7.9, 10mM KCl, 0.5mM Spermidine, 0.1% TritonX-100, 20% glycerol, freshly added protease inhibitors). Nuclei were pelleted at 600g for 3 minutes at 4°C, the supernatant was discard and nuclei were resuspended in 600μL NE buffer. During the cell washes, Concanavalin A beads (Polysciences) were prepared. For 500,000 nuclei 200μL bead slurry was used, for 50,000 nuclei 150μL bead slurry was used, for 5,000 nuclei, 100μL bead slurry was used, for 500 nuclei and 50 nuclei 50μL bead slurry was used, and for 10 nuclei 20μL bead slurry were used. Beads were transferred to a microfuge tube containing 3X volume cold Binding buffer (20mM HEPES-KOH, pH 7.9, 10mM KCl, 1mM CaCl_2_, 1mM MnCl_2_). Beads were washed twice in 1mL cold Binding buffer and resuspended in 300μL binding buffer. Nuclei were added to beads with gentle vortexing and incubated for 10 minutes at room temperature.

*Antibody binding:* After nuclei binding, the supernatant was discarded and bead-bound nuclei were blocked with 1mL cold Blocking buffer (20mM HEPES, pH 7.5, 150mM NaCl, 0.5mM Spermidine, 0.1% BSA, 2mM EDTA, freshly added protease inhibitors) which was added with repeated gentle pipetting and incubated for 5 minutes at room temperature. The supernatant was discarded and nuclei/beads were washed in 1mL cold Wash Buffer (20mM HEPES, pH 7.5, 150mM NaCl, 0.5mM Spermidine, 0.1% BSA, freshly added protease inhibitors) and resuspended in 250μL cold Wash Buffer. Primary antibody was added with gentle vortexing of bead-bound nuclei in 250μL cold Wash Buffer to a final concentration of 1:100. Samples were incubated with rotation at 4°C for 2 hours. The supernatant was discarded and samples were washed twice in 1mL cold Wash Buffer. The supernatant was discarded and samples were resuspended in 250μL cold Wash Buffer.

*Protein A-micrococcal nuclease (pA-MN) binding and cleavage:* pA-MN (kindly provided by Steven Henikoff) was added with gentle vortexing of the nuclei in 250μL cold Wash Buffer to a final concentration of 1:400. Samples were incubated with rotation at 4°C for 1 hour. The supernatant was discarded and samples were washed twice in 1mL cold Wash Buffer. The supernatant was discarded and samples were resuspended in 150μL cold Wash Buffer. Samples were equilibrated to 0°C on ice water for 5-10 minutes. To initiate cleavage, 3μL 100mM CalC_2_ was added during gentle vortexing, samples were flicked quickly to mix and returned to ice water. After 5 minutes of digestion, reactions were stopped with addition of 150μL 2XSTOP buffer (200mM NaCl, 20mM EDTA, 4mM EGTA, 50ug/mL RNaseA, 40ug/mL glycogen, 10pg/mL yeast spike-in DNA). Samples were incubated at 37°C for 20 minutes to digest RNA and release DNA fragments. Samples were centrifuged at 16,000g for 5 minutes and supernatants were transferred to a new microfuge tube while pellets and beads were discarded. Following addition of 3μL 10% SDS and 2.5μL 20mg/mL Proteinase K, samples were mixed by inversion and incubated at 70°C for 10 minutes. DNA was purified using phenol/chloroform/isoamyl alcohol (PCI) extraction followed by chloroform extraction and precipitated with glycogen and ethanol. DNA was pelleted with a high-speed spin at 4°C, washed, air dried for ~5 minutes and resuspended in 36.5μL 0.1XTE.

*Library preparation:* Libraries were prepared using a modification of the Henikoff protocol
(http://blocks.fhcrc.org/steveh/papers/Codomo_Solexa_library_prep_protocol.docx). DNA end-repair, phosphorylation, and A-tailing was performed in a single reaction, as follows. T4 DNA Polymerase (NEB) was diluted 1:20. 5μL 10X T4 DNA ligase buffer (NEB), 2.5μL 10mM dNTPs, 1.25μL 10mM ATP, 3.13μL 40% PEG4000, 0.63μL T4 PNK (NEB), 0.5μL diluted T4 DNA Polymerase, and 0.5μL Taq polymerase (homemade) was added to 36.5μL of CUT&RUN enriched DNA. Samples were incubated at 12°C for 15 minutes, 37°C for 15 minutes, followed by 72°C for 20 minutes in a thermocycler. Samples were put on ice immediately and the following adapter ligation reaction was performed. PE Illumina adapters with inline barcodes were used for these experiments. 55μL of 1X Quick ligase buffer (NEB), 5μL Quick ligase (NEB), and 5μL of 1.5μM adapter mix was added to 50μL of A-tailed DNA and samples were incubated at 20°C for 15 minutes in a thermocycler. Immediately following adapter ligation, samples were purified using Ampure XP beads (Beckman Coulter). Beads were warmed to room temperature during adapter ligation and 38μL well-mixed beads were added to libraries. Samples were mixed thoroughly and incubated for 15 minutes at room temperature. Following solution clearing on a magnetic rack, supernatants were discarded and beads were washed two times with 200μL 80% EtOH. Samples were briefly spun, residual liquid was discarded, and beads were allowed to air dry for ~5 minutes. DNA was eluted from beads by resuspending beads in 30μL 10mM Tris-HCL pH 8.0 and incubation at room temperature for 2 minutes. Following solution clearing on a magnetic rack, 27.5μL DNA was transferred to a 0.2mL PCR tube and libraries were amplified by PCR as follows. 10μL 5X KAPA buffer, 1.5μL 10mM dNTPs, 5μL 20μM PE PCR 1.0, 20μM PE PCR 2.0, 1μL KAPA HotStart HiFi polymerase (KAPA) was added to 27.5μL of library DNA. The following PCR program was used: 98°C 45 seconds, 98°C 15 seconds, 60°C 10 seconds, steps two and three were repeated the specified number of times, followed by 72°C 1 minute. For DNA isolated from 500,000 or 50,000 cells 14 cycles was used, for DNA isolated from 5,000 cells 16 cycles was used, for DNA isolated from 500 cells 17 cycles was used, for DNA isolated from 50 cells 19 cycles was used, for DNA isolated from 10 cells 21 cycles was used. The number of PCR cycles was originally determined using qPCR following 5 cycles of initial amplification using a procedure previously described for ATAC-seq library preparation (Buenrostro et al., 2013; 2015a). Following library amplification, samples were loaded on an agarose gel and DNA corresponding to 150-700 bp was gel extracted using Qiagen gel extraction buffer and Econospin columns. The size distribution of libraries was determined using a Fragment analyzer, and libraries were sequenced on an Illumina NextSeq500.

### Single Cell uliCUT&RUN

Single cell uliCUT&RUN samples were prepared as above with the following alterations to the protocol. Single cells were sorted into individual wells of a 96-well plate containing 100μL NE buffer using a BD FACSAria II Cell Sorter. 15μL of Concanavalin A beads were washed twice with Binding buffer, resuspended in 50μL of Binding buffer, and added directly to the wells containing individual cells. During the 10-minute binding incubation, samples were mixed by pipetting and transferred to 1.5mL microfuge tubes. After discarding the cytoplasmic fraction and blocking the sample in Blocking buffer, beads were washed and resuspended in 125μL Wash buffer. Primary antibody was added during gentle vortexing in 125μL wash buffer to a final concentration of 1:100. Following a 2-hour incubation with rotation at 4°C and washing, beads were resuspended in 125μL Wash buffer and pA-MN was added to a final concentration of 1:400, in 125μL Wash buffer during gentle vortexing. Following two washes, beads were resuspended in 150μL Wash buffer and equilibrated to 0°C in ice water for 5-10 minutes. 3μL 100mM CaCl_2_ was added during gentle vortexing and digestion was permitted to proceed for 30 minutes. Chelation of the reaction was performed with 2XSTOP buffer containing only 1pg/mL yeast spike-in DNA. Release of fragments and extraction was performed as above. Library preparation was performed as above with the following changes. Libraries were amplified for 22 cycles and samples were size selected twice by running two successive agarose gels.

### Embryo collection and microinjections

Superovulated FVB females were mated with FVB males and zygotes were collected ~17-19 hours post-hCG. Zygotes were washed in M2 medium with hyaluronidase to remove surrounding cumulus cells, followed by additional 3 washes in M2. Zygote microinjections were performed on Zeiss AxioVert200 microscope using an Eppendorf Femtojet microinjector. Approximately 5pL of esiRNAs against GFP, Brg1 and Nanog were injected per zygote, at the concentration of 0.5μg/μL. Zygotes were subsequently placed in KSOM media in 5% CO_2_, 5% O_2_ incubator and cultured until the blastocyst stage. Blastocysts were washed twice in M2 and the zona pellucida was removed by acid Tyrode solution, followed by two additional M2 washes. Individual blastocysts were then pipetted into NE buffer.

### RT-qPCR

RNA was isolated from the cytosolic fraction of blastocysts using Agencourt RNAClean XP beads (Beckman Coulter). Isolated RNA was used to synthesize cDNA with a mixture of oligo-dT and random hexamers (Promega). cDNA was used in quantitative PCR reactions with *Smarca4* or *Nanog* specific primers and a FAST SYBR mix (KAPA Biosystems) on an Eppendorf Realplex.

### Immunofluorescence

Zygotes were harvested and microinjected as above and cultured to the blastocyst stage. The zona pellucida was removed and embryos were fixed in 4% PFA. Immunostaining was performed as previously described (http://www.ijdb.ehu.es/web/paper.php?doi=052073mt). Antibodies against NANOG and BRG1 were used at 1:200 dilution. The secondary antibody used was AlexaFluor 546 goat anti-rabbit IgG (Molecular Probes) at 1:500 dilution. Stained blastocysts were either mounted on coverslips or in drops of Vectashield mounting medium with DAPI to retain their three-dimensional structure. Microscopy was performed on AxioObserver.Z1 /7 microscope using 63X/1.4 NA oil objective (coverslips) or 40x/1.3 NA oil objective (drops).

### Blastocyst uliCUT&RUN

Blastocyst uliCUT&RUN samples were prepared as above with the following alterations. Harvested blastocysts were washed twice in M2 and the zona pellucida was removed by Acid Tyrode solution, followed by two additional M2 washes. Individual blastocysts were then pipetted into 300μL NE buffer, centrifuged for 2 minutes at 600g, and incubated on ice for 10 minutes. 20μL of Concanavalin A beads were washed twice with Binding buffer, resuspended in 150μL of Binding buffer, and added directly to tubes. Samples were incubated at room temperature for 10 minutes. Blocking, incubation with primary antibody, and incubation with pA-MN was performed as above. Following two washes, beads were resuspended in 150μL Wash buffer and equilibrated to 0°C in ice water for 5-10 minutes. 3μL 100mM CaCl_2_ was added during gentle vortexing and digestion was permitted to proceed for 30 minutes. Chelation of the reaction was performed with 2XSTOP buffer. Release of fragments and extraction was performed as above. Library preparation was performed as above with the following changes. Libraries were amplified for 18 cycles and samples were size selected twice by running two successive agarose gels.

### uliCUT&RUN data analysis

Paired-end reads were trimmed to 25 bases, barcodes were removed, and reads were then aligned to mm10 using Bowtie2 with the parameter -X 1000. Duplicates were then removed using Picard (http://broadinstitute.github.io/picard/). Reads with low quality score (MAPQ < 10) were removed. Reads were separated into the following size classes: <120bp for TF occupancy and 150-500bp for nucleosome occupancy. These reads were processed in HOMER (Heinz et al., 2010). Genome browser tracks were generated from mapped reads using the “makeUCSCfile” command. Mapped reads were aligned over specific regions using the “annotatePeaks” command to make 20 bp bins over regions of interest and sum the reads within each bin. Peaks were called using the “findPeaks” command, and peaks were compared using the “mergePeaks” command. Motifs were identified using the “findMotifs” command. Published ChIP-seq datasets compared to CUT&RUN data were: CTCF (GSE11431); H3K4me3 (GSE31039); OCT4 (GSE11724); SOX2 (GSE11724); NANOG (GSE11724); BRG1 (GSE14344); H3K27ac (GSE31039); and EZH2 (GSE49435). These datasets were aligned, converted to mm10 using LiftOver, and processed in HOMER. Peaks were called using the “findPeaks” command.

To test for enrichment of low and high cell number uliCUT&RUN samples over background, we summed the normalized reads within 100 bp surrounding each binding site and performed a Kolmogorov-Smirnov test (K-S test) of whether the distributions of peak enrichment values were significantly different for factor-specific antibodies and no antibody controls. To test for differences in NANOG enrichment in control and *Smarca4* KD blastocysts, we subtracted background enrichment from peak enrichment in each KD. Average peak height for each embryo was calculated by summing the average reads within the 100 bp surrounding NANOG peaks. Average background was calculated by summing the reads from -2000 to -1000 bp and dividing by 10 to generate an average background read density per 100 bp. Background was subtracted from peak for each embryo for the data plotted in Figure 4E. A Mann-Whitney U test was used to assess whether differences in *EGFP* and *Smarca4* KD embryos were significant.

### ATAC-seq data analysis

ATAC-seq datasets analyzed were from H1 ESC 50,000 cells (GSE85330) and 27 randomly selected H1 ESC single cells (GSE65360). Paired-end reads were trimmed to 24 bases and reads were then aligned to hg19 using Bowtie2 with the parameter -X 2000. Duplicates were removed using Picard (http://broadinstitute.github.io/picard/). Reads with low quality score (MAPQ < 10) and reads mapping to the mitochondrial genome (chrM) were removed. Reads were separated into size classes as described (Buenrostro et al., 2013) and only nucleosome free reads (less than 100 bp) were used for subsequent analyses. Peaks were called using the “findPeaks” command. Overlapping peaks were identified using the “mergePeaks” command.

---

**Supplemental Figure 1.**
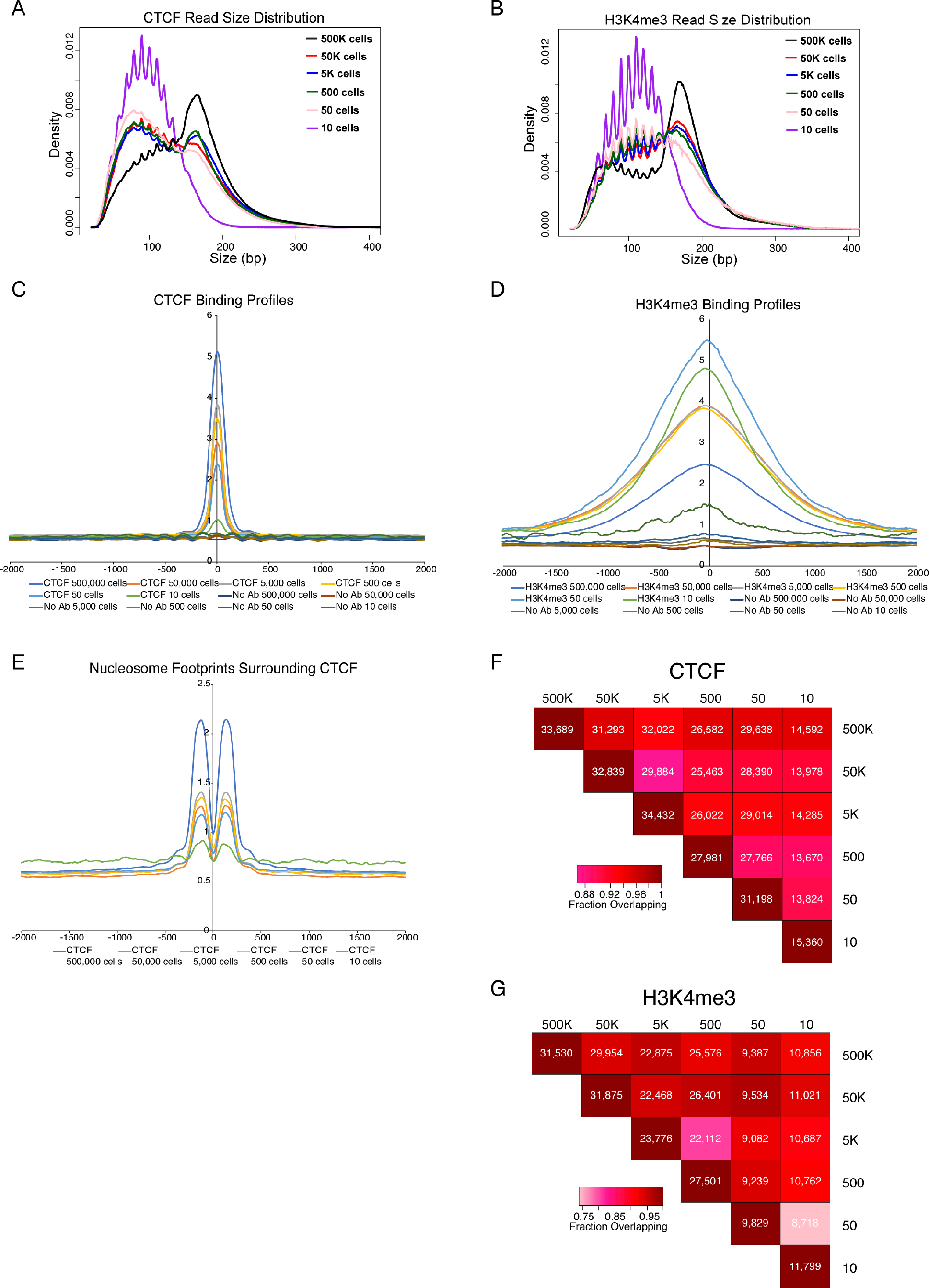
Profiles of CTCF and H3K4me3. **(A-B)** Density plots of the read size distributions of CTCF (A) or H3K4me3 (B) uliCUT&RUN from various numbers of cells. **(C)** Average enrichment surrounding previously published CTCF binding sites of 1-120 bp uliCUT&RUN reads from CTCF and no antibody control experiments from indicated numbers of cells. **(D)** Average enrichment surrounding previously published H3K4me3 binding sites of 150-500 bp uliCUT&RUN reads from H3K4me3 and no antibody control experiments. **(E)** 150-500 bp uliCUT&RUN reads from CTCF uliCUT&RUN libraries shown in (C), revealing well-positioned nucleosomes flanking CTCF binding sites. **(F)** Overlap of CTCF binding sites identified by uliCUT&RUN for various cell numbers. For each uliCUT&RUN experiment, the number of peaks overlapping with published CTCF binding sites was determined as described in the text. The color of each box reflects the fraction of binding sites in the lower cell number library that are shared with the higher cell number for each comparison. The number listed in each box is the number of overlapping peaks. **(G)** Overlap of H3K4me3 binding sites. Data are depicted as in (F).

**Supplemental Figure 2.**
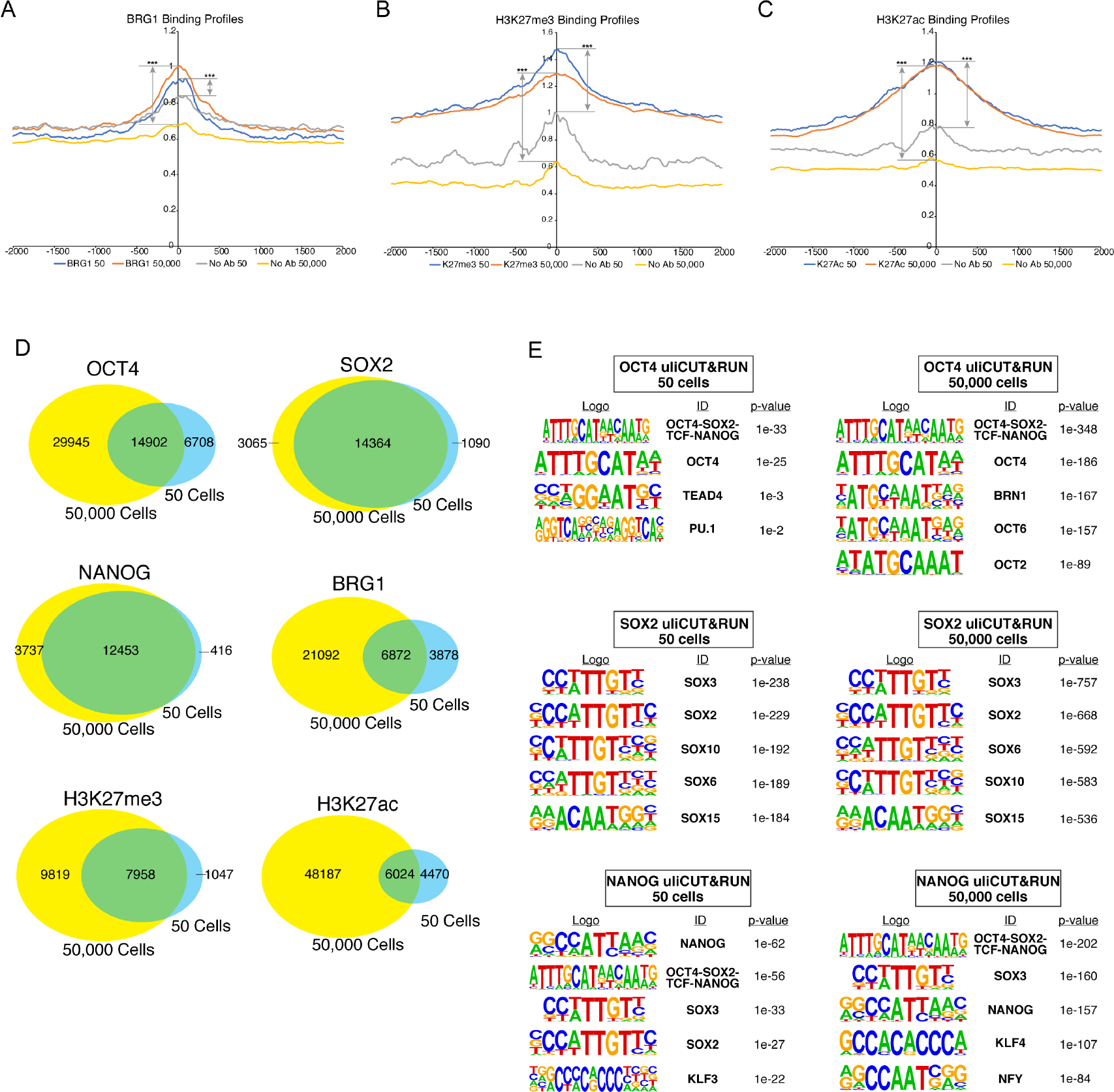
Comparisons of factor enrichment from low and high cell number experiments. **(A-C)** Average enrichment of uliCUT&RUN reads surrounding ChIP-seq identified binding sites of each factor from indicated numbers of cells. Enrichment of 1-120 bp reads shown for BRG1 (A) and 150-500 bp reads shown for H3K27me3 (B) and H3K27ac (C). Significance of enrichment of each factor over no antibody controls was tested using a K-S test (see methods). ***p<2.2×10^−16^. **(D)** Shown are Venn diagrams depicting the overlap of peaks from 50 and 50,000 cell uliCUT&RUN experiments. Peaks for each factor at each cell number corresponding to known binding sites were identified for 50,000 cells (yellow) and 50 cells (blue) their degree of overlap (green) was determined. **(E)** Motifs enriched in peaks of enrichment from 50 and 50,000 cell uliCUT&RUN experiments mapping OCT4, SOX2, and NANOG. Shown are the top five most significant motifs enriched within each sample, including the DNA logo, its corresponding TF, and its p-value. See methods for details. Note, for the OCT4 50 cell uliCUT&RUN experiment, only four motifs had significant p-values and are shown.

**Supplemental Figure 3.**
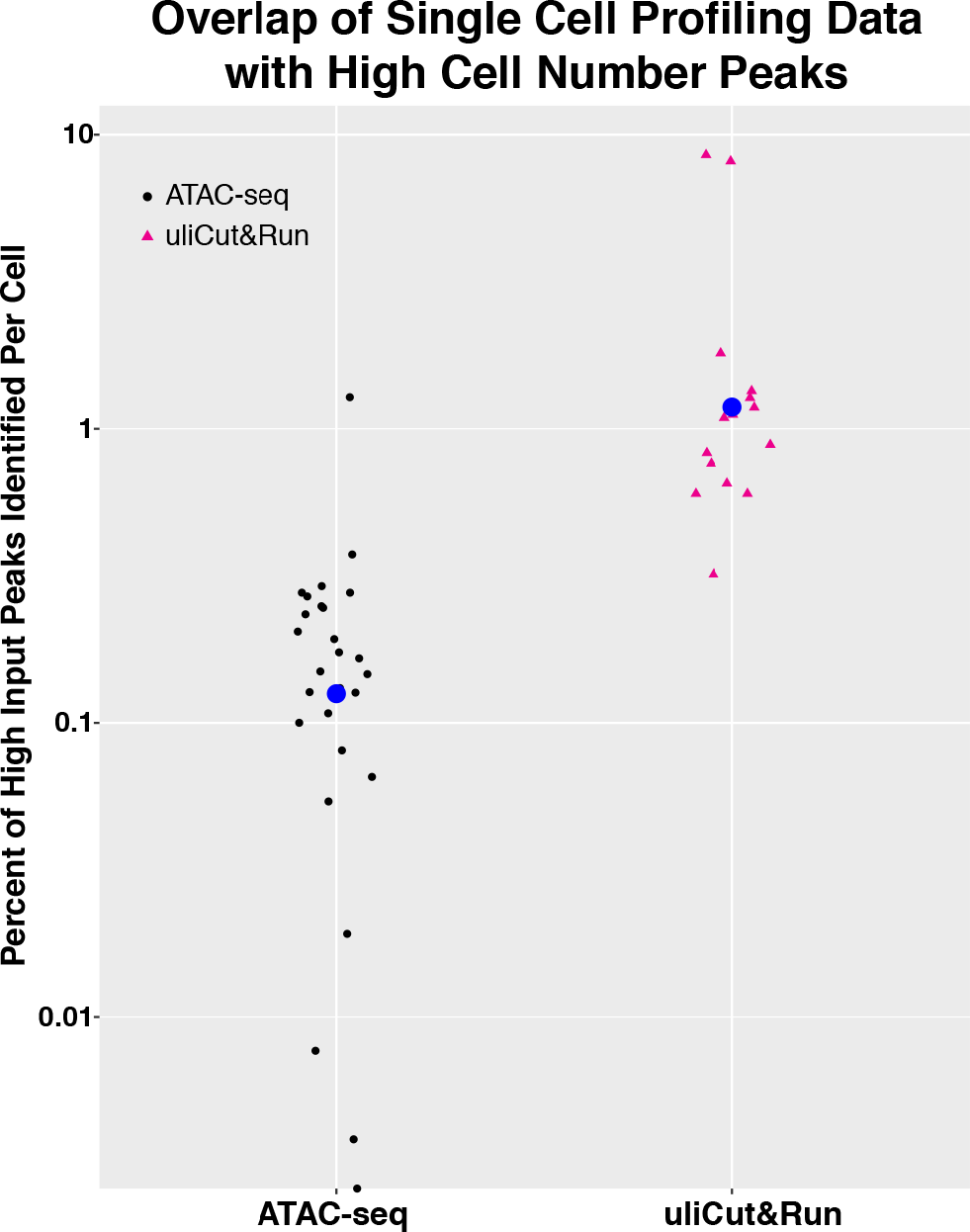
Peak identification by two single cell epigenomic profiling methods. Plotted are the percent of high input ATAC-seq (left) and uliCUT&RUN (right) peaks identified in single cell experiments. Blue dots denote the averages. The percent of ATAC-seq peaks from high input H1 hESCs (GSE85330) identified in each of 27 randomly selected H1 single cell ATAC-seq libraries (GSE65360) are shown. Similarly, the fraction of high input ChIP-seq peaks for CTCF, SOX2, and NANOG identified by single cell uliCUT&RUN for the same three factors are plotted. The percentage of uliCUT&RUN peaks identified for each cell can be found in Supplementary Table 1.

**Supplemental Figure 4.**
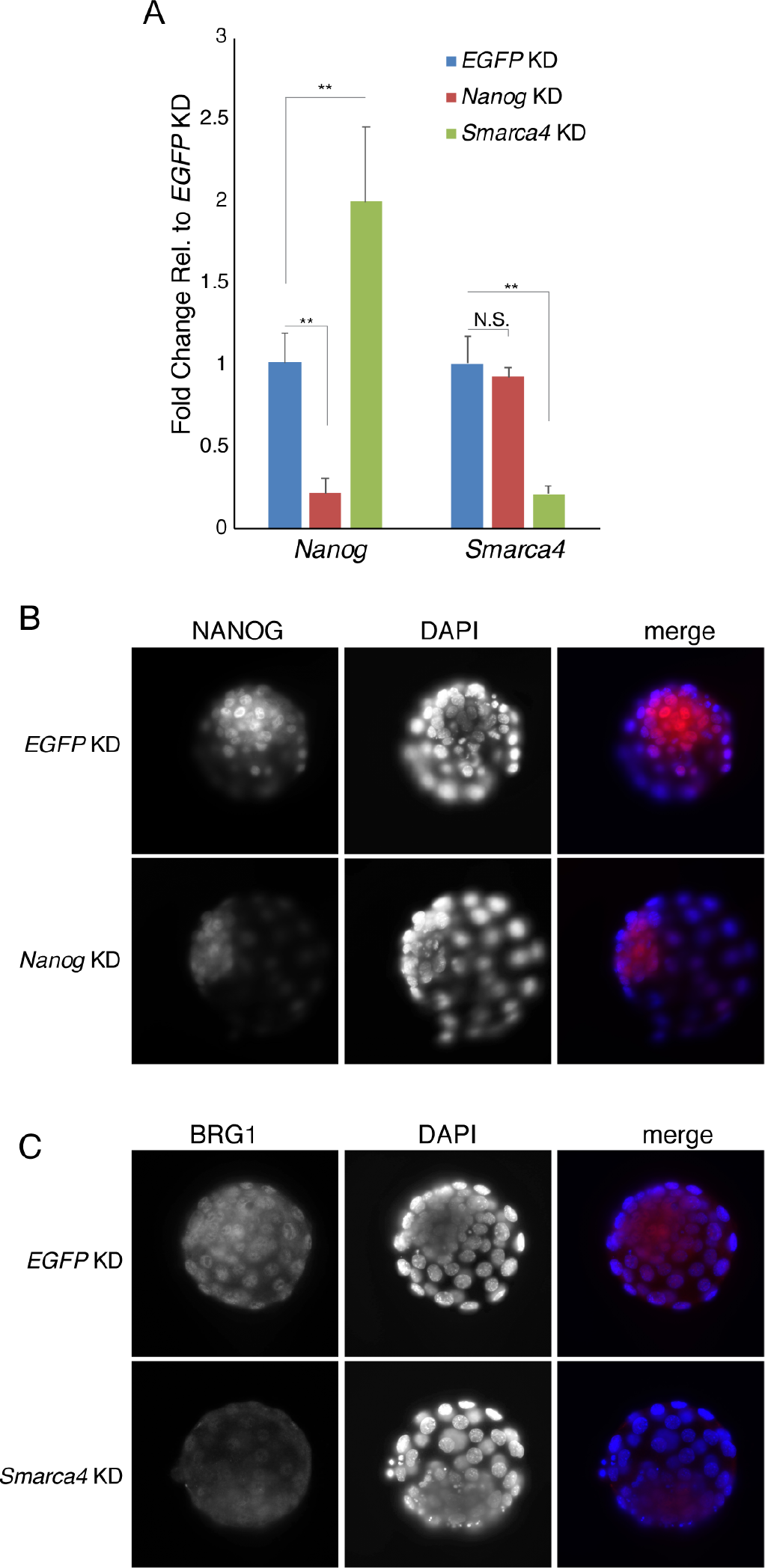
Efficient knockdown (KD) in mouse blastocysts upon injection of validated esiRNAs. **(A)** RT-qPCR of cytoplasmic RNAs from embryos injected with indicated esiRNAs at the one cell stage and cultured to blastocysts. **P<0.01, two-tailed Student’s t-test. N.S., not significant. Error bars represent one standard deviation. **(B-C)** Immunofluorescence for NANOG (B) or BRG1 (C) in blastocyst stage embryos upon KD of *Nanog* or *Smarca4*, respectively by injection of esiRNAs at the one cell stage, as above.

